# HREM, RNAseq and cell-cycle analyses reveal the role of the G2/M-regulatory protein, Wee1, on the survivability of chicken embryos during diapause

**DOI:** 10.1101/2022.02.23.481599

**Authors:** Narayan Pokhrel, Olga Genin, Dalit Sela-Donenfeld, Yuval Cinnamon

**Affiliations:** Agriculture Research Organization, The Volcani Center, Department of Poultry and Aquaculture Science, Rishon LeTsiyon, Israel; Koret School of Veterinary Medicine, The Robert H. Smith Faculty of Agriculture, Food and Environment, The Hebrew University of Jerusalem, Rehovot, Israel

**Keywords:** Chicken embryonic diapause, Cell cycle, G2/M transition, Chicken embryonic blastoderm, Wee1, High Resolution Episcopic Microscopy (HREM), RNAseq, Mitosis

## Abstract

Avian blastoderm can enter into diapause when kept at low temperatures, and successfully resume development (SRD) when re-incubated in body-temperature. These abilities, which are largely affected by the temperature and duration of the diapause, are poorly understood at the cellular and molecular level. To determine how temperature affects embryonic morphology during diapause, High-Resolution Episcopic Microscopy (HREM) analysis was utilized. While blastoderms diapausing at 12°C for 28 days presented typical cytoarchitecture, similar to non-diapaused embryos, at 18°C much thicker blastoderms with higher cell-number were observed. RNAseq was conducted to discover the genes underlying these phenotypes, revealing differentially-expressed cell-cycle regulatory genes. Amongst them, *Wee1*, a negative-regulator of G2/M transition, was highly expressed at 12°C compared to 18°C. This finding suggested that cells at 12°C are arrested at the G2/M phase, as supported by bromodeoxyuridine incorporation (BrdU) assay and phosho-histone-H3 (pH3) immuno-staining. Inhibition of Wee1 during diapause at 12°C resulted in cell-cycle progression beyond the G2/M and augmented tissue volume, resembling the morphology of 18°C-diapaused embryos. These findings suggest that diapause at low temperatures leads to *Wee1* upregulation which arrests the cell-cycle at the G2/M phase, promoting the perseverance of embryonic cytoarchitecture and future SRD. In contrast, *Wee1* is not up-regulated during diapause at higher temperature, leading to continuous proliferation and maladaptive morphology associated with poor survivability. Combining HREM-based analysis with RNAseq and molecular manipulations, we present a novel mechanism that regulates the ability of diapaused-avian embryos to maintain their cytoarchitecture via cell-cycle arrest, which enables their SRD.

## 1. Introduction

The avian embryo at oviposition, termed blastoderm, corresponds to the blastocyst stage in mammalian embryos [1]. However, unlike mammalian embryos, growth and development of the avian embryo can be arrested for a prolong duration at low temperature prior to gastrulation, in a phenomenon known as diapause [2]. In nature, the ability to diapause allows for obtaining synchronous hatching of eggs laid in a clutch. The poultry industry utilizes the diapause phenomenon to store eggs prior to incubation [3,4]. The ability to diapause for long duration, and the capacity to successfully resume development (SRD) thereafter, highly depends on the temperature at which the embryos are kept during diapause. We have recently demonstrated that embryos stored at 12°C have a greater chance to SRD following prolonged diapause than embryos stored at 18°C [3]. The reasons for this variability could be explained by activation or inhibition of different molecular and cellular processes at the corresponding diapause temperatures.

At oviposition, the blastoderm consists of 60,000-130,000 cells distributed in an outer ring termed the Area Opaca (AO), which forms extra embryonic structures, and an inner disk known as the Area Pellucida (AP), which is composed of an epithelium layer, termed epiblast, that gives rise to the embryo proper [3,5–8]. From the epiblast, single cells normally detach and undergo poly-ingression in a ventral direction. These polyingressing cells contribute to form the hypoblast, the second layer of blastoderm. Thus, blastoderms are undergoing continuous cytoarchitectural changes during development, which result in morphological remodeling. We have previously shown that blastoderms that diapause for a prolong time at 18°C undergo overt morphological changes together with an increase in tissue size, as compared to embryos kept at 12°C, yet the latter have higher mitotic index [3]. These counter-intuitive findings led us to suggest that the higher mitotic index at 12°C manifests a mitotic arrest, rather than rapid proliferation. Moreover, we have demonstrated during diapause at 12°C embryos present more healthy-looking and viable cells than at 18°C [3]. The sufficient count of healthy living cells in an appropriate cytoarchitectural organization and differentiation state are prerequisites for embryos to SRD following diapause [3,4,9]. Nevertheless, it is yet unknown whether cell-cycle regulatory mechanisms, such as cell cycle progression play a central role in regulating the embryo’s characteristics including cell survival and morphological remodeling during diapause.

The cell-cycle is regulated by multiple regulators at different phases and check-points of the cell cycle. At the G1 phase, the tumor suppressor retinoblastoma protein (pRb) interacts with the E2F transcription factor to inhibit cell cycle progression [10]. Phosphorylation of pRB, releases E2F which promotes the expression of Cyclin genes, such as cyclinD and cyclinE, which drives the cells to enter into the S phase [11–13]. By late S-G2 phases CyclinA and cyclinB expression increases and degrades during mitotic exit [13–15]. At the G1/S transition cyclinE interacts with cyclin dependent kinase 2 (CDK2), whereas during S-G2-M phases, cyclinA interacts with CDK2/1, and cyclinB interacts with CDK1 [16]. Importantly, several checkpoint kinases, such as Wee1, CHK1, CHK2, and Rad53 have been found to inhibit CDK’s activities directly or indirectly during the G1/S/G2 phases, thereby preventing cell cycle progression [17–19]. For instance, at the G2-M checkpoint, the kinase Wee1 directly phosphorylates CDK1 and prevent the cell from entering mitosis [20]. At cell cycle arrest, the checkpoint kinases are also indispensable for survival of early stage mouse embryos, at the blastocysts stage [21]. Importantly, inhibition of checkpoint kinase activity of Wee1 in mouse embryos by addition of the Wee1 inhibitor MK1775 [22] dramatically shortens G2 phase to ∼30 minutes, which is only 25% of the expected length of G2 phase [23]. This accelerated mitotic entry is accompanied by compromised genomic integrity in blastocyst stage mouse embryos, as lack of Wee1 activity can lead to DNA damage, chromosomal instability, and aneuploidy [24]. Wee1 plays a similar role in maintaining genomic integrity in the mammary gland and acts as a haploid tumor suppressor since a mutant mammary gland develops tumors [25]. Crucial for maintaining genome integrity, Wee1 is a potential target for developing therapies for treating cancer in human patients, [26], and given the high evolutionary conservation, the chick embryo can serve as a model organism for efficacy studies, including toxicity, genome integrity, teratogenicity, and cell survival.

Based on our recent findings, which showed that blastoderms which diapause at 12°C or 18°C have a different morphology and ability to undergo cell division [3], in this study we aimed at determining the possible link between cell-cycle regulation, blastoderm cytoarchitecture and the ability to SRD following diapause. Using high resolution episcopic microscopy (HREM), qualitative and quantitative analyses were performed to characterize in detail the difference in cell proliferation, poly-ingression, cell death and tissue remodeling during diapause at 12°C or 18°C. Moreover, a comparative RNAseq analysis has been performed, revealing various cell cycle regulatory genes that display differential expression between the two diapause temperatures. Interestingly, the expression of the G2/M transition regulator kinase *Wee1*, was significantly increased at 12°C-diapaused embryos in comparison to the 18°C-ones. Further investigation of the role of Wee1 demonstrates that during diapause at lower temperatures, cell cycle is predominantly arrested at the G2/M transition via Wee1, which is associated with better maintenance of cell viability, blastoderm cytoarchitecture and the ability to SRD. These findings are the first to uncover molecular processes that are involved in diapause phenomenon in avian.

## 2. Materials and Methods

### 2.1. Egg collection and isolation of blastoderm

Freshly-laid Ross (308) broiler eggs from young flock age (28-45 weeks) were collected from a commercial farm and stored at 18°C or 12°C up to 28 d in a cooled incubator (SN 265959, VELP SCIENTIFICA, Italy). The temperature and relative humidity of cooled incubator was continuously monitored throughout the experiment. Following diapause, blastoderms were isolated as previously described [27]. Briefly, the egg shell was carefully cracked, the albumin was discarded and the yolk with the embryo was gently placed in a beaker containing phosphate buffered saline (PBS, REF-BP507/1LD, Hylabs, Israel). The vitelline membrane, surrounding the underlying embryo was incised with scissor and gently separated from the remaining yolk. The separated vitelline membrane with the embryo was transferred to a petri dish containing PBS. The embryo was then cleaned by removing the adherent yolk from the embryo surface by gently washing with drops of PBS. The isolated blastoderms were either placed in RNA save solution (Cat no. 01-891-1A, Biological Industries, Israel) for RNA stabilization or fixed in 4% paraformaldehyde (PFA) in PBS.

### 2.2. HREM imaging

Embryos were fixed overnight in 4% PFA, washed with PBS and dehydrated in series of methanol concentration (25%, 50%, 75%, and 100%, each for 20 min). Following dehydration the embryos were impregnated with 50% methanol and 50% JB4-dye mix solution, for overnight on a rotating shaker at 4°C. JB4 dye mix contains solution A (Cat no. 14270-01, Electron Microscopy Sciences, USA), 12.5 mg/ml catalyst (Cat no. - 14270-06, Electron Microscopy Sciences, USA), 2.75 mg/ml Eosin B (Cat no. −861006-10G, Sigma, USA) and 0.5 mg/ml Acridine orange (Cat no.-A6014-10G, Sigma, USA), dissolved for overnight at 4°C and then filtered.

The 50% methanol and 50% JB4-dye mix solution was replaced with 100% JB4-dye mix solution, and impregnation continued for 24h at 4°C. For curing the resin at mounting, 30µl/ml of solution B (Cat no.-14270-04, Electron Microscopy Sciences, USA) was added to the JB4-dye mix, following by curing the resin overnight at 4°C in a plastic embedding mold (Cat no.-15899-50, Polysciences Europe GmbH, Germany). Polymerized sample blocks were further hardened by overnight baking at 70°C.

Cooled hardened sample blocks were mounted in the HREM machine (Serial no. - 007, Indigo scientific, UK), and block-surface images were serially taken every 2.76µm section, throughout the entire sample in 2700 × 1800 image pixel resolution. Depending on the experimental design, selected plastic sections obtained during HREM sectioning were preserved by placing them on DDW drops on histological slides, and let them dry at room temperature. Images of plastic section were captured using microscope (Serial no. – 469580, model – DM2000LED, Leica, Germany). Image of reference 1000µm graticule (EMS, USA) was taken for each block sectioned, to allow for accurate size measurements.

All the images acquired from HREM imaging were processed, 3D reconstructed and analyzed as previously described [3,27,28] using Photoshop (Adobe Photoshop CS. (2004), Berkeley, CA: Peachpit Press), Fiji software [29] and Amira software (FEI, Hillsboro, Oregon USA) for 3-D modeling of blastoderm morphology. Briefly, the HREM acquired serial images were inverted, processed, and converted to 8 bit using Photoshop (Adobe Photoshop CS. (2004), Berkeley, CA: Peachpit Press) and serial images were stacked, rotated and cropped using Fiji software [29]. The staked images were then accessed for 3D reconstruction using Amira software. Specific region of embryos was selected and compared by quantifying their volume using the Amira software segmentation tool. The data were exported for further statistical analysis. At least 4 embryos were used for each HREM image analysis.

### 2.3. RNA-Seq analysis

RNA seq gene expression profiling of embryos was done in four different storage conditions (0d-unstored, 7d/18°C, 28d/18°C and 28d/12°C). Each groups included 4 biological repeats. Embryos were isolated as previously described [27] to obtain 16 individual blastoderms. RNA was extracted from cells with RNeasy micro kit (Qiagen, cat no. 74004) using the Qiacube automated system (Qiagen). RNAseq libraries (cat no. E7760, NEBNext UltraII Directional RNA Library Prep Kit for Illumina, USA) from 16 samples were produced according to manufacture protocol using 200 ng total RNA. mRNAs pull-up was performed using Magnetic Isolation Module (cat no. E7490, NEB, USA). All 16 libraries were mixed into a single tube with equal molarity. The RNAseq was generated on Illumina NextSeq500, 75 cycles, high-output mode (cat no. FC-404-2005, Illumina, USA). Quality control of single–end reads were assessed using Fastqc (v0.11.5). Reads were then aligned to Chicken reference genome and annotation file (Gallus_gallus-5.0 and Gallus_gallus.Gallus_gallus-5.0.93.gtf downloaded from ENSEMBL) using STAR aligner (STAR_2.6.0a). The number of reads per gene was counted using Htseq (0.9.1). The differential gene expression (DEG) analysis and bioinformatics analysis of DEGs were done as described previously [30–32]. DEGs between 28d/18°C and 28d/12°C were analyzed using WebGestalt [30]. Over-representation (enrichment) analysis was used for pathway analysis with the KEGG functional database [33,34]. Heat map of enriched gene sets was generated using clustvis web based tool [35].

### 2.4. Real time PCR analysis

Three isolated embryos were pooled for RNA extraction. Subsequently, cDNA was prepared using Promega kit (REF 017319, Promega, USA). Primers were designed according to the sequence information from the NCBI database using Primer3 Input (version V. 0.4.0) software [36]. qRT-PCR was performed in a final reaction volume of 10µl with the SYBR Green PCR Master Mix Kit (REF-4309155, Applied biosystems by Thermo Fisher Scientific, UK) in the applied biosystems Real-Time PCR Detection System (SN 2720011007, Applied biosystems stepOnePlus Real-Time PCR System, Singapore), using the following program: 95°C/20 seconds and 40 cycles of 95°C/1 second, and 60°C/20 seconds. All reactions were performed in duplicates for each sample, and GAPDH was used as a reference gene for normalization of gene expression levels. The relative gene expression values were calculated using the 2^-ΔΔct^ method. Comparison of gene expression levels were analyzed by one-way anova statistical tool. The list of primers that were used to analyze the expression of genes are given in Supplementary table 1.

### 2.5. BrdU incorporation assay in-ovo in embryos

Fresh, fresh +6 hr incubation for positive control, 7d/18°C, 7d/12°C, 28d/18°C and 28d/12°C stored embryos were treated with 0.2 mg/ml BrdU (REF B9285-1G, Sigma, USA) in PBS, by injecting 100 µl in volume to at least 4 embryos through a shell window. Following injection, the egg shell window was sealed using Leukoplast (REF 72668-02, BSN medical GmbH, Germany), and placed back to their respective condition for 6 hrs. The embryos were isolated, fixed in 4% PFA overnight at 4°C and dehydrated in methanol (series of steps: 25%, 50%, 75% and 100%). Dehydrated samples were stored at −20°C in 100% methanol. Whole mount embryo samples underwent DNA denaturation steps by incubating them in 50% formadide (CAT No 00068023G500, Bio-Lab, Israel), in 4XSSC in DDW, for 2 hr at 65°C under slow agitation. Subsequently, samples were rinsed in 2XSSC for 15 min and incubated in 2N HCL at 37°C for 30 min, followed by neutralization in 0.1 M sodium borate (pH 8.5) for 10 min. The samples were then washed 6 times, 15 min each, in TBS (0.15 M Nacl and 0.1 M Tris-HCL, pH 7.5) and blocked for 1-hr with 3% Normal Bovine serum in TBS containing 3% triton-X-100 (TBST, CAS-9002-93-1, Sigma, USA). Following blocking, embryos were incubated with rat anti-BrdU IgG2a (1:200; REF-OBT0030G, Serotec, UK) for overnight at 4°C, washed 3 times in TBS for 15 min each, at room temperature, rinsed once in TBST for 15 min, and incubated for 2 hr with F(ab)2 donkey anti-rat IgG-Cy3 antibody (1:200; REF-712-166-153, Jackson ImmunoResearch Laboratories, USA). Embryos were washed in TBS stained with 4’,6-diamidino-2-phenylindole (DAPI) (1:1000, REF-D9542, Sigma, USA) for 30 min at RT. Stained embryos were mounted between two cover-slips using fluorescence mounting medium (REF-9990402, Immu-mount, Thermo Scientific, USA), and scanned using a confocal microscope (magnification, X10 and X40; Leica TCS SPE, Germany). The DAPI stained nuclei and BrdU positive cells were counted manually to determine percentage of BrdU incorporating cells.

### 2.6. Immunohistochemistry

Embryos were isolated and fixed in 4% PFA overnight, washed with PBS, permeabilized with 0.2% triton in PBS (PBST), blocked in 10% NGS in PBST at 4°C for 3 hr and immunohistochemistry steps were carried out by incubating the samples with primary anti-pH3 antibody (diluted 1:300 in blocking buffer, REF 05-817R, clone 63-1C-8, recombinant rabbit monoclonal antibody, Millipore, USA) for overnight at 4°C as previously described [37–39]. The primary antibody was washed in PBST and incubated for overnight at 4°C with secondary Alexa fluor 594 conjugated anti-rabbit IgG antibody, diluted 1:300 in blocking buffer (REF A11012, Invitrogen, USA). Embryos were washed in PBS, stained with DAPI, mounted and scanned as described above. pH3 positive cells and DAPI stained nuclei were counted manually to calculate percentage of pH3 positive cells. Moreover, the distribution of M phase in diapaused embryos was calculated by further dividing the M phase into sub-phases, M1 to M7 based on pH3 staining that showed chromatin condensation and chromosomal rearrangement and alignment [40–43]. At least four embryos were analyzed per experimental group.

### 2.7. Embryo treatment with wee1 inhibitor-MK1775

Firstly, 15% pluronic gel (P2443-250G, Sigma, USA) was prepared in PBS at 4°C and mixed with MK-1775 to a final concentration of 500 nm (HY-10993, MedChemExpress LLC, USA) [17,22,44,45]. Through a hole in eggshell to the air sac, 250 µl of MK-1775 containing gel, or gel only as control, was injected in area near to embryo. The window in the eggshell was sealed using Leukoplast (REF 72668-02, BSN medical GmbH, Germany) and embryos were stored back at 12°C for either 24 h or 7d, depending on the experimental design. In particular, embryos diapaused for 6 d at 12°C were treated with MK-1775 and stored at 12°C for an additional 24 h, whereas fresh embryos treated with MK-1775 were stored for 7 d at 12°C. At least 4 embryos from each group were then isolated as described above, fixed in 4% PFA for at least 24 h and accessed for immunohistochemistry for pH3 staining and Terminal deoxynucleotidyl transferase dUTP nick end labeling (TUNEL) assay.

### 2.8. TUNEL assay

Whole mount embryos were immunostained with anti-pH3, followed by dehydration as described above. The embryos were cleared in xylene and processed for paraffin 7µm sections as described before [3] using microtome (RM2035, Leica, Germany). Following deparaffinization and rehydration, the sections were immersed in 0.85% NaCl for 5 min, washed in PBS for 5 min, post fixed for 15 min in 4% PFA for 15 min, and washed in PBS. The sections were treated with PBS containing 2% Triton X-100 for 5 min and washed in PBS. TUNEL staining was done using DeadEnd Fluorometric TUNEL detection system (REF-G3250, Promega, USA). For positive control, sections were treated with 25µl/ml DNAase I (REF-M0303S, BioLabs, New England) in DNase I buffer (pH 7.9 40mM Tris-HCl, 10mM NaCl, 6mM MgCl_2_ and 10mM CaCl_2_) for 10 min, fixed in 4% PFA, and washed in DDW. The TUNEL staining was performed according to manufacturer instructions of TUNEL detection kit, then the samples were stained with DAPI, washed, and mounted with coverslip and mounting media (REF-9990402, Immu-mount, Thermo Scientific, USA). The intensity of TUNEL-positive cells and pH3-positive cells were quantified as previously described [46–48].

### 2.9. Statistical tests

Data were analyzed using the statistical tests, t-test, one-way ANOVA and two-way ANOVA. Data were presented as mean ± SEM. Statistical tests were performed using JMP (John’s Macintosh Project) software (JMP, 189-2007, SAS Institute Inc., Cary, NC, USA) at significance level (P< 0.05).

## 3. Results

### 3.1. The cellular morphology of the blastoderm during diapause

The cytoarchitectural structure of prolong diapaused-embryos was investigated by 3-dimensional (3D) HREM imaging [3,28,49,50]. To qualitatively analyze the overall thickness of the blastoderms, the 3D reconstructed images (Figure 1A-C) were subjected to a transparency test against a checker-board background (Figure 1D-F, upper panel). The transparency test showed that embryos diapaused for 28 d at 18°C (Figure 1F, upper panel) are more opaque than those diapaused at 12°C or than freshly-laid embryos (Figure 1D,E, upper panels). To confirm that the transparency test emanates for the difference in embryonic thickness, the HREM images (Figure 1D-F, middle panel) and plastic sections (Figure 1D-F, lower panel) of the blastodermal tissue were analyzed at a high-resolution. Images acquired from the plastic sections showed that the polyingressing cells in fresh embryos migrated ventrally (Figure 1D, lower panel), whereas embryos diapaused for 28 d at 18°C were thicker with notable recesses at the dorsal side of the epiblast (Figure 1F, lower panel, blue arrowhead; also showed in Figure S1). Notably such recesses were not apparent in embryos diapaused at 12°C (Figure 2E, lower panel).

**Figure 1.**
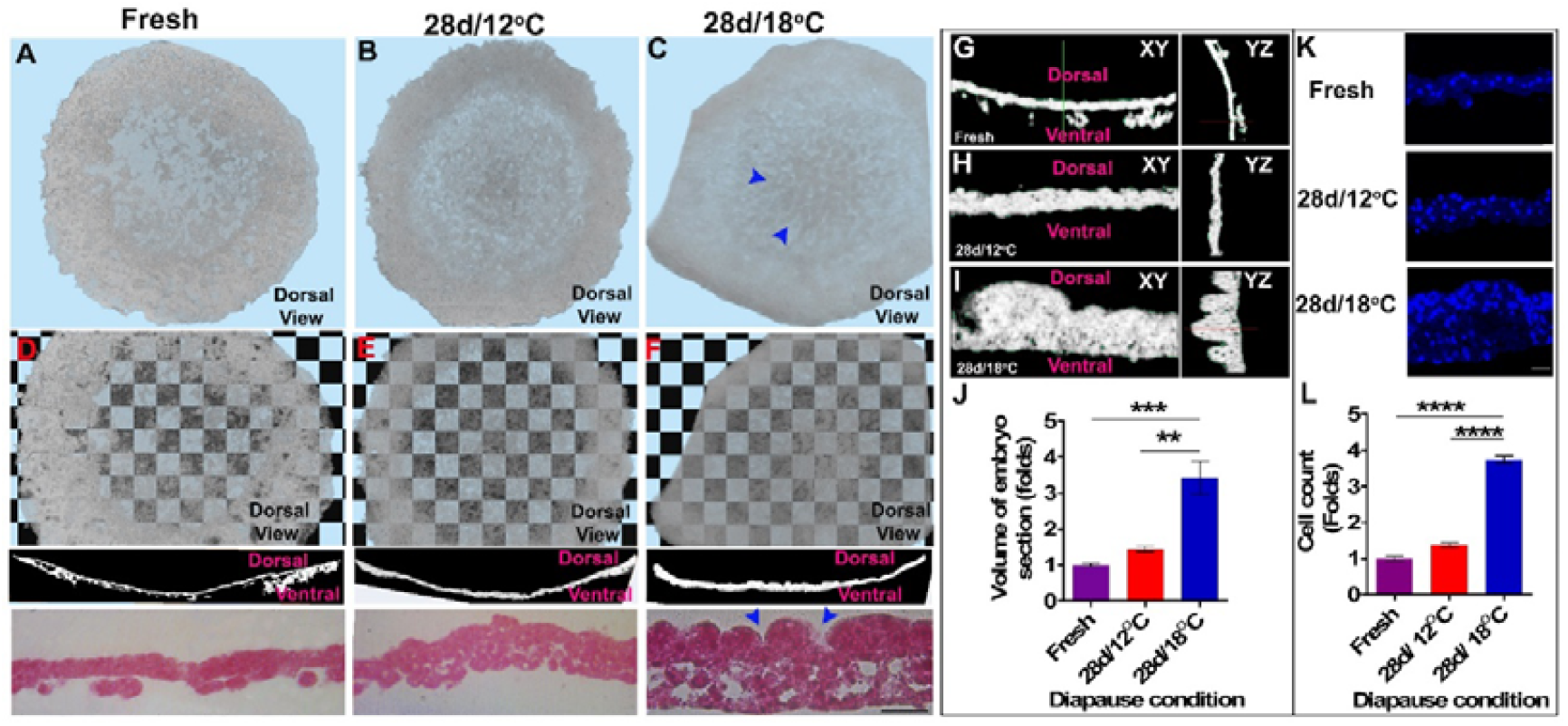
Cellular morphology of non-diapaused and prolong diapaused (28d/12°C and 28d/18°C) blastoderm. Blastoderm were sectioned by using High Resolution Episcopic Microscopy (HREM) and processed by AMIRA software to obtain 3-dimentional (D) image. (A to C) Dorsal views of blastoderms are showed. (A) Fresh, (B) 28d/12°C, (C) 28d/18°C. Tissue remodeling in terms of recess formation is absent in 28d/12°C while it is present in 28d/18°C embryos (arrowhead). (D to F) Checkerboard transparency test of fresh (D) and prolong diapaused blastoderms (28d/12°C (E) and 28d/18°C (F), upper panel), their corresponding HREM sections (D and F, middle panel), and plastic sections (D and F, lower panel, bar size 100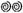m). Plastic section of 28d/18°C embryos reveals that epiblast contract to form recess (C, blue arrowhead) and the ventral polyingressing cells contribute in thickening of dorsal epiblast layer (F, lower panel). (G to I) XY and YZ planar views of fresh (G) and prolong diapaused embryo (28d/12°C, H, and 28d/18°C, I) showing their dorsal and ventral layers. J. Volumetric analysis of fresh and prolong diapaused embryos. Embryos have bigger tissue volume following prolong diapause for 28 d at 18°C than fresh and 28d/12°C (one way anova; **p=0.0021; ***p=0.0007). K. Embryos were sectioned in 7 microns thickness and stained with DAPI. Tissue sections were imaged using confocal laser microscopy to view nuclear staining of blastoderms. Bar size 20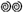m. L. Cell count analysis. Embryos consist of more number of cells following diapause at 28d/18oC, compared with fresh and 28d/12°C (One-way ANOVA; ****p<0.0001)..

**Figure 2.**
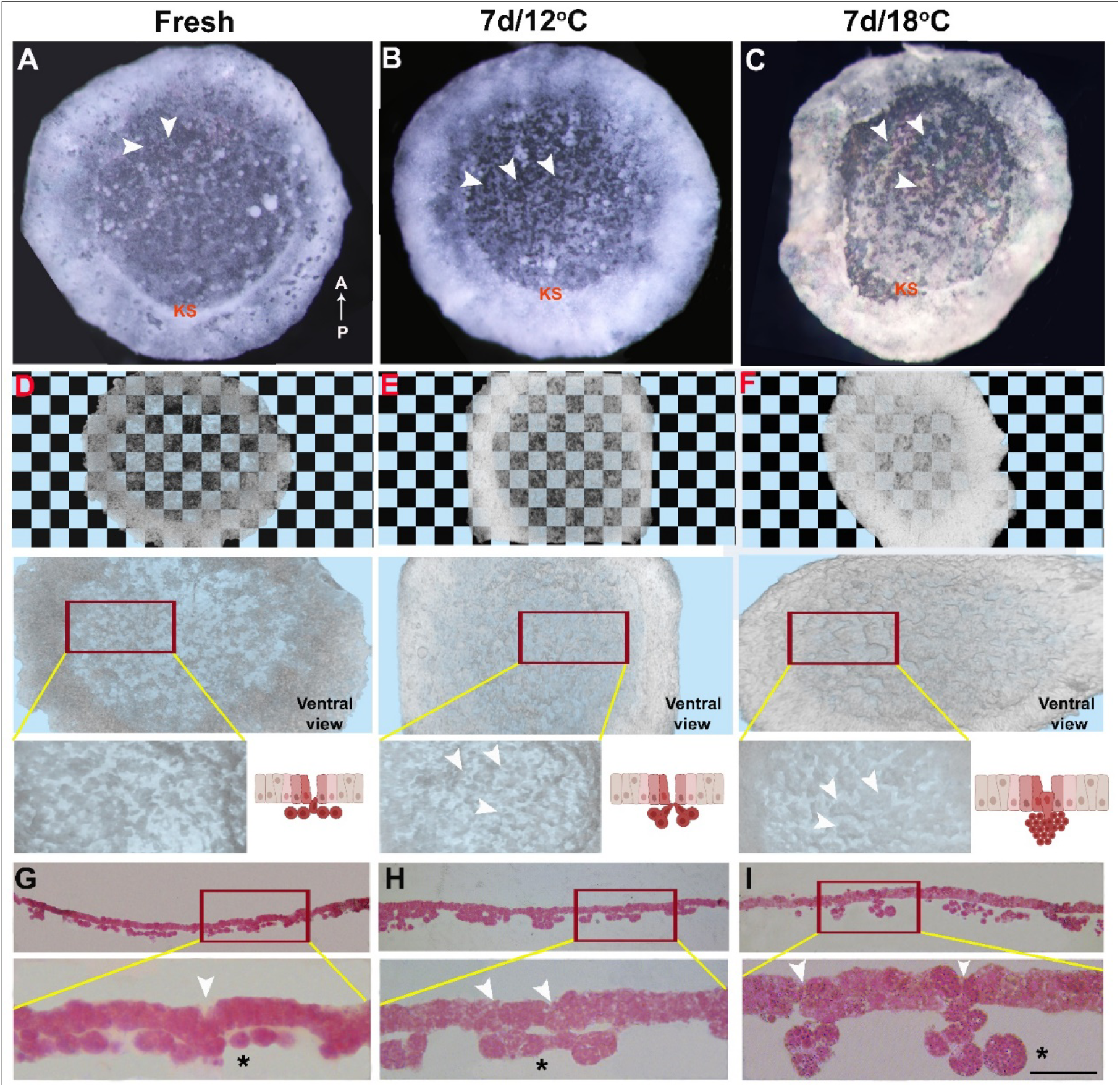
Cellular morphology of blastoderm reveals cytoarchitectural changes already within 7 d of embryo diapause at higher temperature. **(A)** Fresh. **(B)** 7d/12°C. **(C)** 7d/18°C. Polyingressing cells are present in anterior region (arrowhead) and Koller’s sickle (KS) marks the posterior region of blastoderm. **(D-F)** HREM images showing dorsal view of blastoderm that underwent checkerboard transparency test. **(D)** Fresh, **(E)** 7d/12°C, **(F)** 7d/18°C. Embryos are more opaque in 28d/18°C condition than in fresh and 28d/12°C. Ventral view of same blastoderm **(D-F**, middle panel) and their polyingressing cells are showed in high magnification HREM images **(D-F**, lower panel, arrow-head) and represented in schematic diagram **(D-F**, lower panel). Cells that undergo polyingression dorso-ventrally from epiblast region have directional cell movement as is observed in fresh (D, lower panel) and 7d/12°C embryos **(E**, lower panel). While already following 7 d of diapause at 18°C, there is disruption in directional cell movement **(F**, lower panel). These differences can be appreciated in plastic sections of the blastoderm obtained following HREM imaging **(G-I**, mag-X20). **(G-I**, lower panel) High magnification images of plastic section (X40). Plastic section shows that blastoderm undergo cellular changes already following 7 d of diapause at 18°C, as revealed by the formation of aggregates of clustered polyingressing cells **(I**, lower panel, *). Bar size 100 μm.

In addition, using multi planar view (MPV) and segmentation tool enabled the visualization of the XY and YZ planes of the blastoderm (Figure 1G-I) and volume quantification (Figure 1J). Quantification of the unit volume of the epiblast region by segmentation tool showed a significant increase in tissue volume of blastoderms which diapaused at 18°C compared to 12°C or to non-diapaused embryos (Figure 1J, **p=0.0021, ***p=0.0007, respectively). To investigate whether the thickening of blastoderm tissues after prolonged diapause at 18°C was due to cell hypertrophy or to cell proliferation, we further sectioned embryos and stained their nuclei with DAPI (Figure 1K), to allow for nuclei count. Quantifying cell number per unit area (Figure 1L), our results showed a significant increase in cell number in the 28d/18°C group compared with the fresh and 28d/12°C groups (Figure 1L, ****p<0.0001), suggesting that cell proliferation majorly accounts for the thicker tissue in embryos diapaused at 18°C.

To identify at what time point the cellular changes are initiated, we compared freshly-laid embryos (Figure 2 A,D,G) with embryos diapaused for 7 d at 12°C (Figure 2B,E,H) or 18°C (Figure 2C,F,I), using HREM sectioning. The 3D images were used for transparency test (Figure 2D-F, upper panel), which demonstrated that already after 7 d at 18°C, the embryos are more opaque compared with the fresh and 12°C groups. In addition, a higher magnification view of the ventral surface (Figure 2D-F, middle and lower panels) revealed that the polyingressing cells at the 7 d/18°C group tend to form large clusters, rather than forming a linear thin layer of the hypoblast. This was further validated by corresponding images from the plastic tissue sections obtained during the HREM process (Figure 2G-I). These findings highlight the aggregation of polyingressing cells after a short diapause at 18°C, which is not observed in 7d/12°C-embryos or in freshly laid embryos (Figure 2G-I, upper and lower panels, asterisk denotes the polyingressing cells).

To quantify the differences between the polyingressing cell clusters between the blastoderms, the volume of cell clusters was individually measured using the 3D segmentation tool (Figure 3). This approach enables to discriminate the polyingressing cells from the entire overlying blastodermal cells. This was done by fixing either freshly-laid embryos (Figure 3A,D,G,J) or those diapaused for 7 d at 12°C (Figure 3B,E,H,K) and at 18°C (Figure 3C,F,I,L) and sectioning them using HREM. To precisely determine the boundaries of the polyingressing cell clusters, multi planar view was used (Figure 3G-I) and the volume of isosurface models (Figure 3J-L) was calculated using Amira software (Figure 3M). The results show that while the size of the polyingressing cell clusters in freshly-laid embryos and embryos diapaused for 7 d at 12°C are similar (p=0.1152), polyingressing clusters in embryos diapaused for 7 d at 18°C are significantly larger (p< 0.0001).

**Figure 3.**
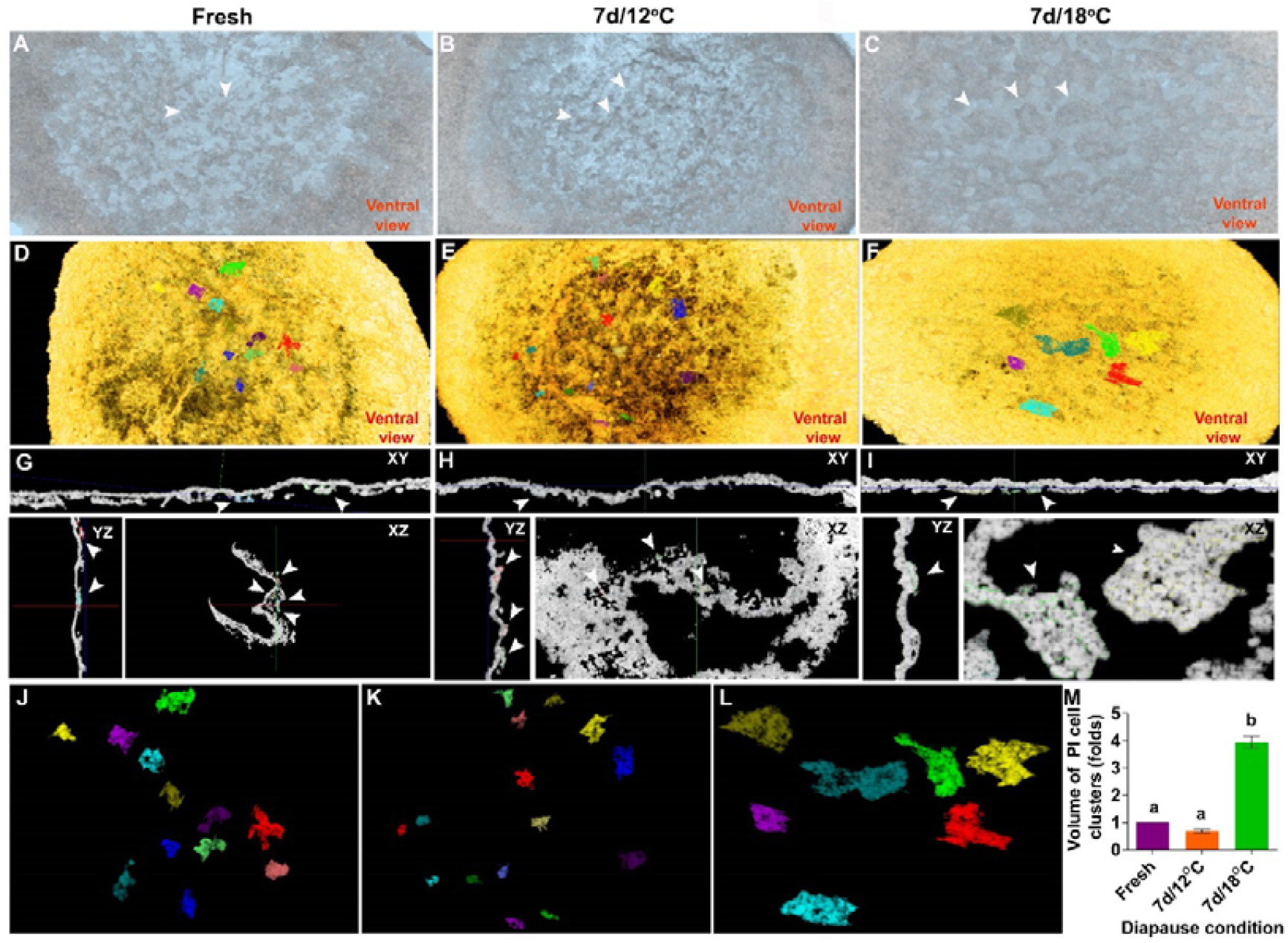
Quantification of volume of polyingressed cells of blastoderm. **(A** to **C)** 3-D image of blastoderm showing their dorsal view. **(A)** Fresh, **(B)** 7d/12°C, **(C)** 7 d/18°C. (**D** to **F**) Polyingressing cells of blastoderm represented in different colors. **(D)**. Fresh, **(E)** 7d/12°C, **(F)** 7d/18°C. **(G** to **I)** Planar views of blastoderm showing polyingressing cells in different colors. **(J** to **L)** polyingressing I cells of blastoderm from fresh, 7d/12°C and 7d/18°C condition. **(M)** Quantification of volume of polyingressing cells (relative in folds). Volume of polyingressing cells is bigger at 7d/18°C condition, compared with fresh and 7d/12°C. Different connecting letters represent that the group are significantly different (One-way ANOVA; a vs. b: p<0.0001).

Altogether, these results indicate that diapausing at 18°C leads to cytoarchitectural changes in the blastoderms that are not found in non-diapaused embryos or in ones diapaused at 12°C. This difference, which is apparent already after 7 d, involves a significant increase in cell cluster formation, thickening of the blastoderm, formation of recesses in the epiblast, and increased cell count.

### 3.2. Transcriptome profiling reveals distinct regulation in cell-cycle in diapaused embryos at 18°C, and 12°C

The major cytoarchitecture changes and the increased cell count found in embryos diapaused at 18°C, suggests that molecular mechanisms regulating the cell-cycle may be involved in these phenomena. To identify these mechanisms, RNAseq analysis was done on 4 groups of embryo: *(i)* freshly-laid embryos which serve as a reference control, *(ii)* embryos diapaused for 7 d at 18°C (7d/18°C), *(iii)* embryos diapaused for 28 d at 18°C (28d/18°C), and *(iv)* embryos diapaused for 28 d at 12°C (28d/12°C). The differences in molecular pathways between 28d/12°C and 28d/18°C groups were analyzed using enriched Kyoto Encyclopedia of Genes and Genomes (KEGG) pathways (Figure 4A). This revealed enrichment of cytoskeletal genes, remodeling of the adherent junctions, regulation of PLK1 activity at the G2/M transition and cell cycle regulation pathways at 28d/12°C, whereas, protein processing in the endoplasmic reticulum, transcriptional misregulation in cancer and toll-like receptor signaling pathway were enriched at 28d/18°C. Further analysis of the enriched gene sets showed that genes involved in cell cycle progression, including cyclins and CDKs, were expressed in the fresh, 7d/18°C, and 28d/12°C groups, while in the 28d/18°C group, these genes were down-regulated (Figure 4B). Notably, *Wee1*, the G2/M transition regulator kinase was found to be upregulated in 28d/12°C (Figure 4B, see asterisk). Moreover, KEGG pathway analysis revealed enrichment of cell cycle pathway and linkage between genes during each cell cycle transition phase (Figure 4C, see also Figure S2), further indicating for differences in the activity of cell cycle-related genes between embryos at different diapause conditions.

**Figure 4.**
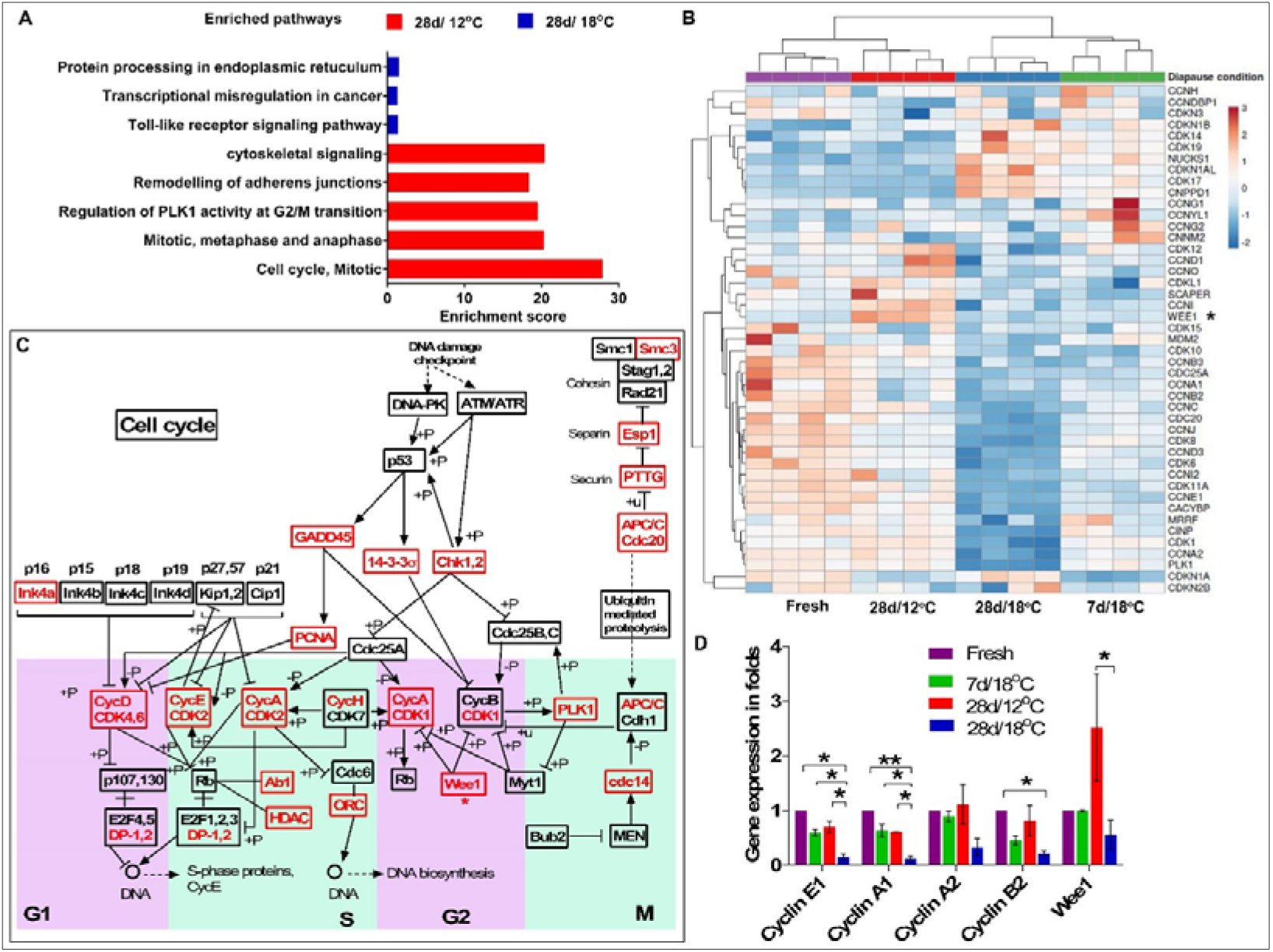
RNAseq analysis and validation. RNAseq analysis was conducted in 4 groups (Fresh, 7d/18°C, 28d/12°C, and 28d/18°C) having 4 replicates each. **(A)** Pathway enrichment analysis between 28d/12°C and 28d/12°C group. **(B)** Heat map of gene sets of embryo in different diapause conditions. **(C)** A schematic representation of enriched cell cycle pathway, as obtained by KEGG pathway analysis of DEGs between 28d/12°C and 28d/12°C group. The enriched gene sets are marked in red. An expanded scheme of enriched pathway is showed in Figure S2. **(D)** The expression of key genes involved in cell cycle were validated by using real-time PCR analysis (One-way ANOVA; *p=0.02;**p=0.0096)..

To examine and validate the expression of enriched cell-cycle related gene-sets in the RNA-seq data, real-time PCR analysis was conducted (Figure 4D). Expression levels of *cyclin E1, cyclin A1/2* and *cyclin B2* were found be down regulated in the 28d/18°C group compared to the other groups (*p=0.02, **p=0.0096), while *Wee1* expression was up-regulated in the 28d/12°C group (*p=0.0485, compared to 28d/18°C group). These results, which confirmed the RNAseq data, indicated deceleration in cell cycling following 28 d of diapause at 18°C. However, the morphological changes observed in 18°C group prompted us to further determine whether there was a shift in cell-cycling from proliferation to deceleration state in the 18°C group, and whether the maintained cyto-architecture at 12°C was associated with safeguarding of cell-cycle progression through higher *Wee1* expression. As a first step to experimentally examine the cell cycle state of embryonic cells, BrdU incorporation assay was performed in order to quantify the number of cells in the S phase of the cell cycle. Different experimental groups were treated with BrdU: 1. As a positive control, fresh embryos were incubated at 37.8° for 6 hours, treated with BrdU and further incubated for 6 hours before fixation. 2. As a starting reference point that refers to embryonic state before diapause, fresh embryos were treated with BrdU for 6 hours at room temperature before fixation. 3. For 7 d time point, embryos were diapaused at 18°C or 12°C for 7 d, then treated with BrdU for 6 hours and fixed, and 4. For 28 d time point, embryos were diapaused at 18°C or 12°C for 28 d, treated with BrdU for 6 hours and fixed (Figure 5A). Following fixation, embryos were stained with anti-BrdU antibody and DAPI to visualize the nuclei, which allowed to calculate the proportion of BrdU positive cells out of the total nuclei count (Figure 5B). Interestingly, our results showed a reduction in BrdU incorporation within embryos diapaused at 18°C already after 7 of diapause, which became statistically significant following 28 d (p=0.001), while at 12°C BrdU incorporation remained stable throughout the 28 d of diapause and was comparable to fresh embryos (Figure 5B-C). These results, which stand in agreement with the RNAseq data, showing stable cyclin’s genes expression over an extended diapause period at 12°C (Figure 4B-D), can suggest a marked deceleration in cell-cycling following extended diapause at 18°C, as opposed to the 12°C group and fresh group. However, the preserved cytoarchitecture and the moderate increase in cell number in prolong diapaused embryos at 12°C, in contrast to the higher cell-count and thicker blastoderms observed at 18°C, argues against such scenario. Hence, an alternative explanation for the results of BrdU incorporation at 12°C is that the cell cycle might be regulated at other transition phases following the S phase, leading to cell cycle arrest at these conditions. Thus, based on RNA-seq analyses (Figure 4B-D), we hypothesized that cell cycle regulation at 12°C may be mediated by the high expression of the negative G2/M transition regulator *Wee1* that prevents the further progression of the cell cycle towards mitosis [20].

**Figure 5.**
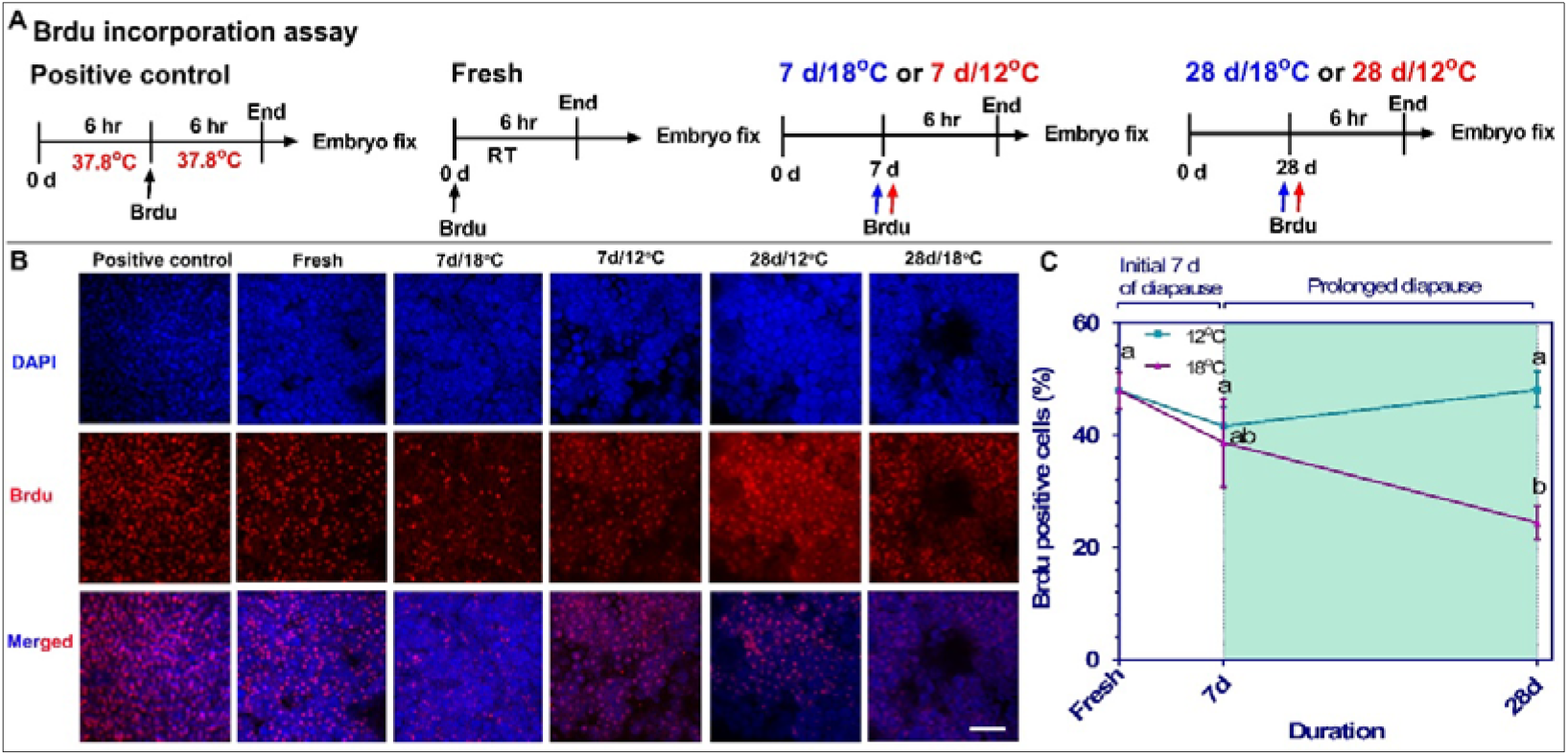
BrdU incorporation assay in blastoderms at different diapause conditions. **(A)** Experimental design. Positive control group consists of 6 hr. incubated embryos that were injected with BrdU for 6 more hr. Likewise, Fresh, 7d/12°C, 7d/18°C, 28d/12°C and 28d/18°C groups were injected with BrdU during 0 d, 7 d, and 28 d of diapause for 6 hr, respectively. Following treatment, embryos were isolated, fixed, immunostained with anti-BrdU antibody and accessed for confocal microscopy. **(B)** BrdU incorporation assay in positive control, fresh, 7d/18°C, 7d/12°C, 28d/12°C and 28d/18°C group. Bars 20 μm. **(C)** Quantification of BrdU positive cells. Prolong diapaused embryonic cells still incorporate BrdU at lower temperature, which is significantly higher than that of cells diapaused at higher temperature (p=0.0159). Different connecting letter refers that the compared groups are significantly different to each other (One-way ANOVA; a vs. b: p=0.001).

### 3.3. Mapping the mitotic phase of diapaused embryos reveals an increase in cell proliferation at higher temperature

The M phase is a complex stage of the cell cycle, involving major reorganization at the cell’s nuclei, cytoskeleton, cytoplasm and membranes [51]. The M phase is subdivided into several stages; prophase, metaphase, anaphase, and telophase, which initiates with chromatin condensation and terminates with the separation into two daughter cells [40,41]. These stages can be further subdivided into more specific events: M1 and M2 stages of the prophase, which refer to the initiation and completion of chromatin condensation, M3 and M4 stages of the pro-metaphase, which refer to the initiation or nearly-completion of chromosome’s segregation, M5 stage of the metaphase, which refers to the align of the chromosomes along the metaphase plate, M6 stage of the anaphase which refers to the separation of the chromosomes into the opposite poles, and M7 stage of the telophase, which refers to the reformation of the nuclei and daughter cells [40,42,43]. As our results demonstrate changes in the proliferation state of embryos stored at different diapause temperatures (Figure 5), as well as difference in the expression levels of the G2/M regulator *Wee1* (Figure 4), here we further examined whether these findings are association with distribution of cells at different sub-phases of the M stage of the cell cycle. Embryos diapaused at 12°C and 18°C, for 7 and 28 d as well as freshly-laid embryos were stained with the M-phase marker pH3 (Figure 6A), and analyzed for their mitotic phase (Figure 6B-D). The mitotic index was calculated based on the chromosomal state at M1-M7 stages (Figure 6C).

**Figure 6.**
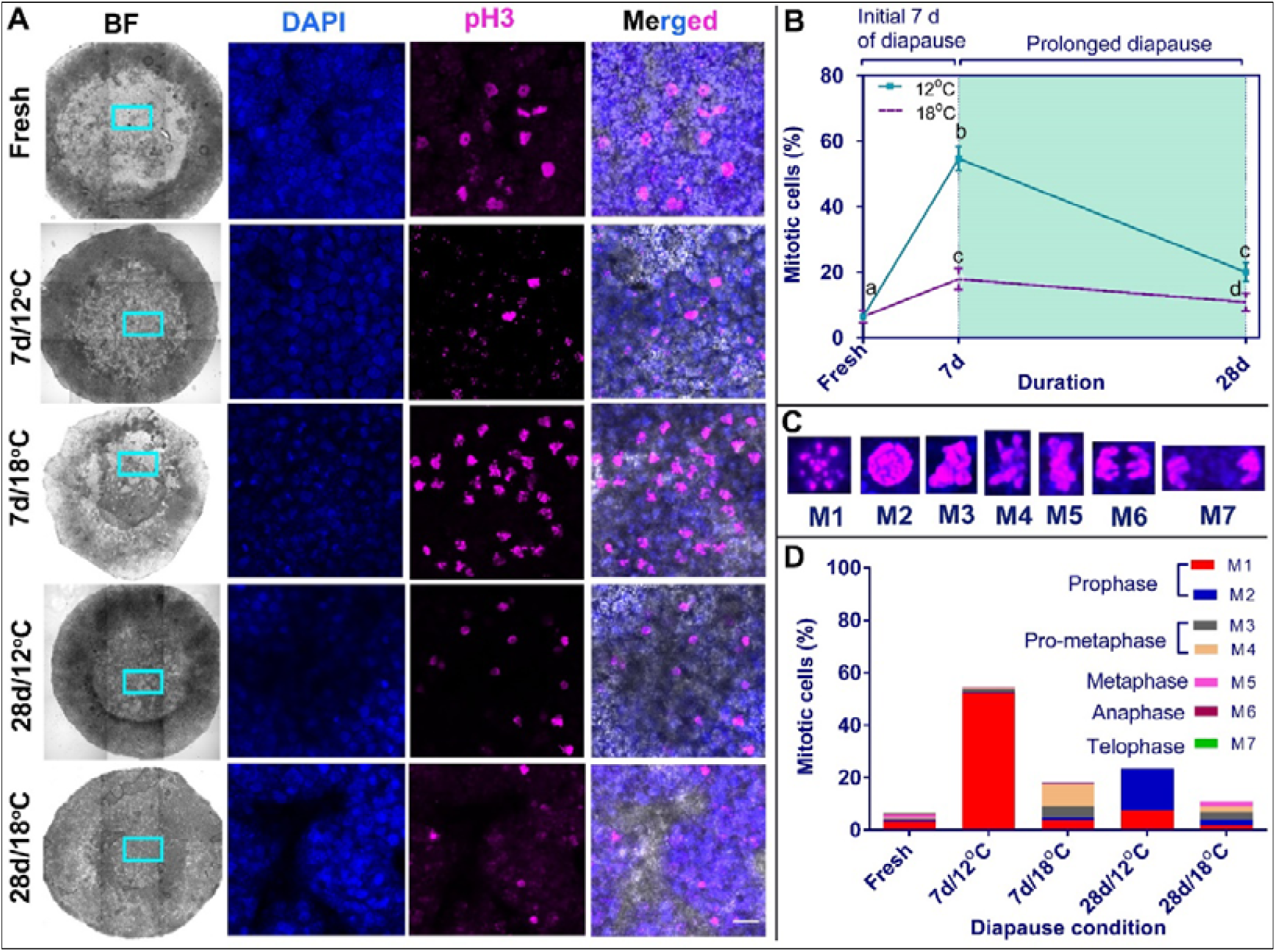
Mapping of mitotic phase of blastoderm in different diapause conditions using pH3 immunostaining. **(A)** Confocal images showing BF, DAPI, pH3 and merged channels of Fresh, 7d/12°C, 7d/18°C, 28d/12°C, and 28d/18°C group. Bars 20 μm. **(B)** Quantification of mitotic cells. Different connecting letter refers that the compared groups are significantly different to each other (One-way ANOVA; b vs. c: p<0.0001; c vs. d: p=0.0028). **(C)** Sub-categorization of mitotic cells based on pH3 staining and nuclear condensation. **(D)** Distribution of mitotic phases in blastoderms that underwent different diapause conditions (fresh, 7d/12°C, 7d/18°C, 28d/12°C, and 28d/18°C).

Calculating the percentage of mitotic/pH3^+^ cells, our results show that at 12°C there are more pH3^+^ mitotic cells than in fresh embryos (Figure 6A,B; fresh vs. 7d, p<0.0001; fresh vs. 28d, p=0.001) and in embryos diapaused at 18°C (Figure 6A,B; during 7d, p<0.0001; during 28 d, p=0.0028). However, quantification of discrete M sub-phases revealed that fresh embryos consisted of cells in all mitotic phases, while embryos diapaused at 12°C were in the prophase stage (p<0.0001), which correspond to the G2/M transition (Figure 6D). In contrast, embryos diapaused at 18°C consisted of cells in pro-metaphase during 7 d (predominant M4 stage, p≤0.002) and in all mitotic sub-phases during 28 d, which distributed similarly along the different stages (Figure 6D). These results suggest that the preserved cytoarchitecture of blastoderms and the higher count of BrdU+ nuclei at the 28d/12°C group (Figure 5) is likely due to the ability of cells to pass the S phase but arrest at the G2/M checkpoint at the prophase, the early-mitosis stage (Figure 6). In contrast, at the 28d/18°C group, the abnormal cytoarchitectural growth may be due to continuous, yet slower, progression of the cell cycle beyond the prophase stage.

### 3.4. Inhibition of Wee1 in embryos diapaused at 12°C results in cell cycle progression beyond prophase

The above results revealed that during diapause at 12°C, the majority of M-phase cells are arrested in prophase, which correlates with the high expression of *Wee1*. Therefore, to determine the role of Wee1 in this G2/M arrest, the small molecule MK-1775, which is a highly specific and potent inhibitor of Wee1 kinase activity [44,45,52,53], was used. Embryos were diapaused at 12°C for 6 d, treated with 500 nm MK-1775 or with PBS (as a control solution) for another 24h, stained for pH3 and quantified for their M1-M7 stage (Figure 7A-D). Our results show that while in the control embryos the majority of the mitotic cells remained at the prophase stage of the M phase (Figure 7C-D, p=0.0079), in MK-1775-treated embryos the cells were found in more advanced stages, from the pro-metaphase M3/M4 stages and onward (Figure 7D, p=0.0111). These results strongly support a regulatory role of Wee1 in preventing cell-cycle progression in embryos diapause at low temperature.

**Figure 7.**
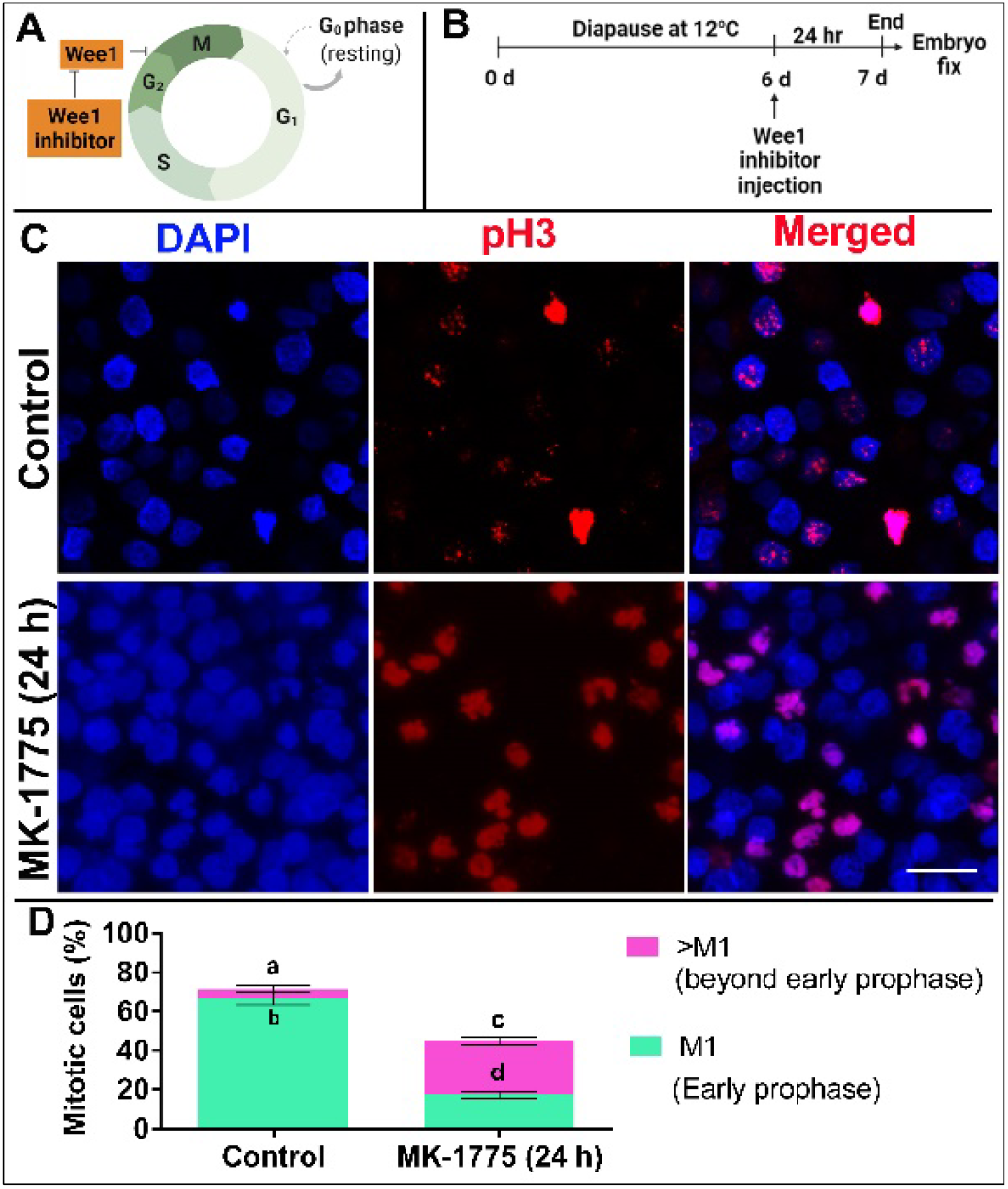
Effect of treatment of Wee-1 kinase inhibitor (MK-1775) on cell cycling of blastoderm. **(A)** Schematic representation of the road map for demonstrating effect of inhibiting wee1 kinase activity on cell-cycling. **(B)** Experimental design. Embryos were subjected to diapause for 6 d at 12°C and subsequently, on d 6, they were injected with Wee1 inhibitor for 24 hr. PBS treated embryos served as control. Following treatment, embryos were isolated, fixed, immunostained with anti-pH3 antibody and accessed for confocal microscopy. **(C)** pH3 immuno-staining of blastoderm that were untreated or treated with MK-1775 for 24 h in 6d/12°C stored embryos. Bars 20 μm. **(D)** Quantification of mitotic cells in blastoderms without or with treatment of MK-1775 for 24 h in 6d/12°C group. Different connecting letter refers that the compared groups are significantly different to each other (Two-way ANOVA; a vs. b: p=0.0079; c vs. d: p=0.011; a vs. c: p=0.0079).

To test whether the activity of Wee1 is coupled with the thinner blastoderm’s phenotype observed in the 12°C-diapausing embryos (Figure 8A), such embryos were treated with PBS (Figure 8B) or MK-1775 (Figure 8C) for 7d and analyzed for their blastoderm cytoarchitecture using HREM sectioning (Figure 8B’,C’,D). Additional embryos were stained for pH3 to quantify their M-phase sub-distribution following 7 d of storage at 12°C (Figure S3). Our results demonstrate that PBS-treated embryos were significantly thinner, with reduced volume (Figure 8B,B’) in comparison to the MK-1775 treated embryos, which demonstrated thicker blastoderms (Figure 8C,C’,D, p<0.0001). Moreover, in control embryos the polyingressing cell clusters were smaller and discreetly arranged (Figure 8C,C’, see also Figure 2B,H), while in the MK-1775-treated embryos the polyingressing cells formed a continuous thick layer, covering the ventral side of the epiblast (Figure 8,C,C’) resembling polyingressing cells of the 18°C–diapaused embryos (Figure 1 and Figure 2). Furthermore, these phenotypes were coupled with an increase in the number of pH3-positive cells in advanced mitotic sub-phases, compared to the PBS-group (Figure S3, p<0.0001). These results suggest that during diapause at 12°C, the increased levels of *Wee1* arrest the cells at the G2/M, thereby inhibiting proliferation and thickening of the blastoderm, while following Wee1 inhibition with MK-1775, the cells progress with the cell cycle, which results in blastoderm thickening, similar to the phenomenon observed during diapause at 18°C when *Wee1* expression levels are low.

**Figure 8.**
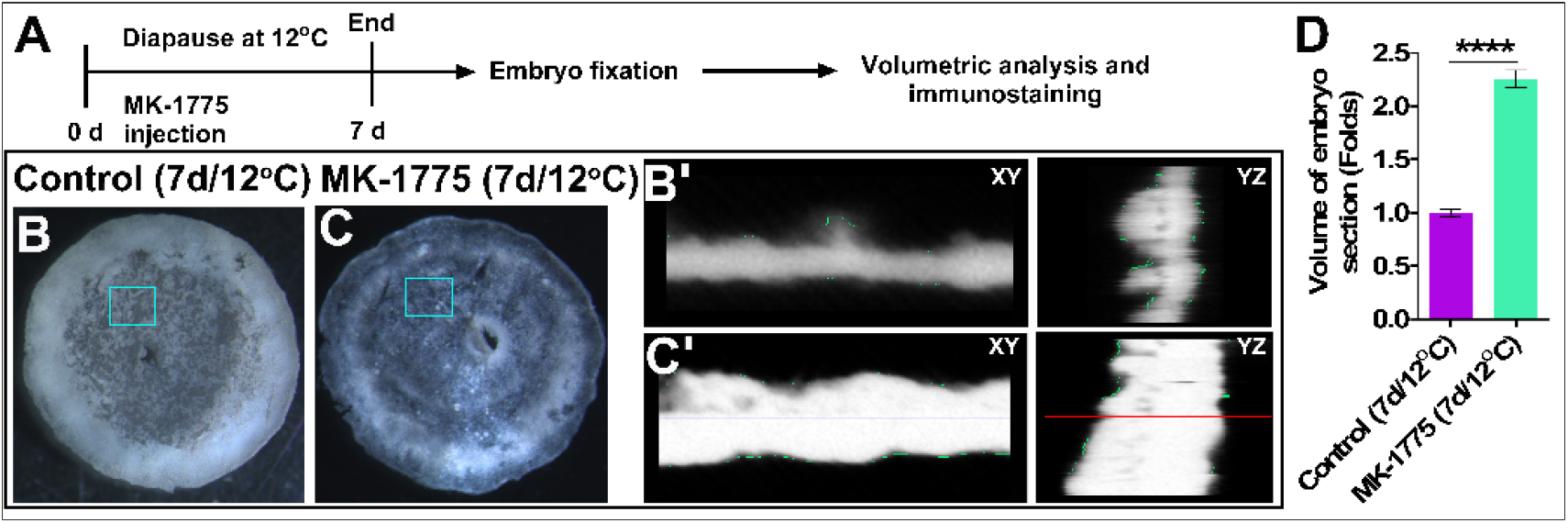
Inhibition of Wee1 kinase activity up to 7 d at 12°C results cellular changes in embryos. **(A)** Experimental design. Fresh embryos were treated with MK-1775 and subsequently, they underwent diapause for 7 d at 12°C. PBS treated embryos under same condition were used as control. Following treatment, embryos were isolated, fixed and subjected to volumetric analysis using HREM 3-D imaging. In a separate experiment, the treated and control embryos were isolated, fixed and immunostained with anti-pH3 antibody and accessed for confocal microscopy (Figure S3). **(B)** Control embryos. (C) MK-1775 treated embryos. While the clusters of polyingressing cells are visible in control embryos, MK-1775 treated embryos undergo cellular changes in terms of hypoblast progression and thickening of tissue. The inset of **B** and **C** is showed in **B**’ and **C**’, which represent the 3-D modeling of embryo section. **(B’)** 3-D images of control embryos and their visualization in XY and YZ planar views. **(C’)** 3-D images of MK-1775 treated embryos and their visualization in XY and YZ planar views. **(D)** Volumetric analysis shows that MK-1775 treated embryos are twice thicker than non-treated control embryos (****p<0.0001).

### 3.5. Increase in staining of cellular death marker, TUNEL upon Wee1 kinase activity inhibition in 12°C-diapaused embryos

As a G2/M checkpoint regulator, Wee1 was found to allow the extension of time spent in repairing DNA damage prior to mitosis [17]. Thus, inability to properly repair DNA damage in cells prior to their progression into the M phase of the cell cycle promotes cell death in other contexts [54,55]. To check whether inhibition of Wee1 kinase activity in 12°C-diapaused embryos induced cell death, such embryos were treated with MK-1775 or PBS and analyzed for apoptotic cell death using the TUNEL assay [56,57], together with their staining for pH3 and DAPI to label the M phase stages (Figure 9A). As a positive control for the TUNEL assay, additional embryos were treated with DNAase to induce DNA damage (Figure 9B). Intensity of TUNEL staining was used to quantify the amount of apoptotic cells (Figure 9C-E) [46,47]. The results show that inhibition of the kinase activity of Wee1 not only increases the embryonic volume (Figure 8D,****p<0.0001) and the number of cells entering M phase (Figure 9E, p=0.0215, see also Figure S3), but also increases the amount of TUNEL-positive cells (Figure 9C-E, p<0.0001). Notably, TUNEL positive cells were also pH3 positive at mitotic phases M4-M5 (Figure 9D). These results further highlight that the G2/M arrest achieved by *Wee1* expression during diapause at 12°C may serve as a survival mechanism for embryos diapaused at low temperatures by preventing accelerated cell proliferation and death and thus, maintaining the cytoarchitectural integrity of the blastoderm.

**Figure 9.**
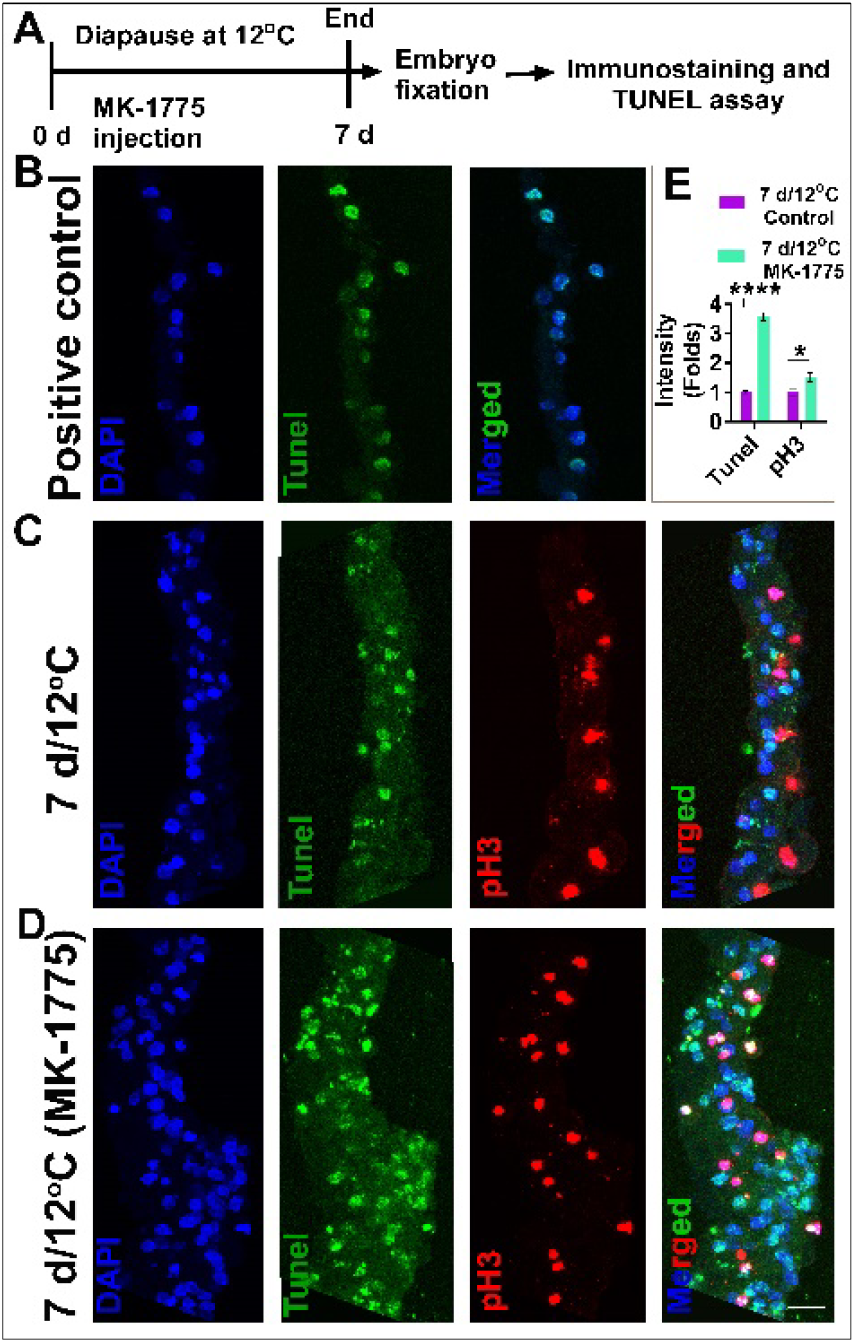
TUNEL assay in diapaused embryos. **(A)** Experimental design. Fresh embryos were treated with MK-1775 and subsequently, they underwent diapause for 7 d at 12°C. PBS treated embryos under same condition were used as control. Following treatment, embryos were isolated, fixed, immunostained with anti-pH3 antibody, sectioned, underwent TUNEL assay in same slide (except for positive control) and accessed for confocal microscopy. Confocal images of all the tissue sections were acquired using the same imaging settings. **(B)** Positive control of TUNEL assay. Sections from fresh embryos were treated with DNAse to induce DNA breaks and accessed for TUNEL assay. **(C)** TUNEL assay and pH3 immunostaining of 7 d/12°C group without or with MK-1775 treatment **(D)**. Bars 20 μm. **(E)** Quantification of fluorescence intensity of TUNEL positive and pH3 positive cells in blastoderms treated with or without MK-1775. t-test, *p=0.0215; ****p<0.0001.

## 4. Discussion

### 4.1. The survivability of embryonic cells during diapause is linked with cell-cycle regulation and maintaining the cytoarchitectural structure

The freshly-laid blastoderm consists of epithelial cells in the AP region that adhere to each other to form a single-layer epithelial sheet [58]. This is promoted by expression of adhesion molecules at the apical and basolateral boundaries such as tight junction protein and integrins [59]. Next, as the blastulation process progresses, some of the AP cells lose their integrity and undergo polyingression in a ventral direction [5]. Our previous study started to characterize the morphological changes that occur during diapause at different time points and temperatures, and demonstrated that at 12 °C, the embryos are more resilient to abnormal morphological remodeling and survive better when they resume their development [3]. Here, we utilized the HREM imaging strategy to fully uncover and quantify the cytoarchitectural differences of blastoderms after prolonged diapause at different temperatures, and by using RNAseq approach, demonstrated for the first time the molecular mechanisms involved for such changes in blastoderm’s cell morphology and cell volume in response to different diapause conditions.

Previously, 2-D imaging strategies were mainly used to study morphology of blastoderms [5], which limited the viewing of the entire embryo in its intact state. Numerous studies have used micro-computed tomography (µCT), optical coherence tomography and scanning electron microscopy (SEM) to obtain 3-D images of bone tissues, post-gastrulating chick embryos, and also the chick blastoderms [60–63], however, the techniques have certain limitation in reconstructing 3-D images of small, thin and delicate tissue like the avian blastoderm. In order to obtain 3-D image of blastoderm, we previously used the HREM image analysis method [27]. This system was found advantageous over conventional blastoderm sectioning methods [5,63,64] and other 3-D imaging methods [60–63] for observing and quantifying various blastodermal components such as AO, AP, polyingressing cells, and hypoblast [27] and also for obtaining high-resolution 3-D images of bigger chick embryos at post-gastrulating stages [28], by utilizing the large data to generate high resolution three-dimensional images [65].

To identify the mechanisms underlying the morphological changes in blastoderms that diapause at different temperature, RNAseq has been performed, and revealing that cell cycle-related genes are distinctly regulated at the two temperatures tested, 12°C or 18°C. KEGG pathway enrichment analysis further suggested that the G2/M transition regulatory gene Wee1 [66] is providing a potential molecular regulator for the G2/M arrest in diapaused embryos at 12°C, which is less active at 18°C, leading to an abnormal progression in the cell cycle and overt changes in morphology. Previously increase in cell number during diapause of mouse embryos has been reported [67]. However, cell cycle arrest during diapause has also been shown in fish embryos [68]. Thus, previous studies have provided evidences that embryos in diapause can either increase the cell number or arrest the cell cycle, which is consistent with the results of this study showing either arrest of the cell cycle or increase in cell number following a small difference in temperature exposure during diapause in chick blastoderms. Cell cycle regulation by Wee1 is not only important for maintaining cytoarchitecture of the blastoderm diapaused at 12°C, but may also be important during the transition from blastulation to gastrulation, because in *Xenopus* it has been shown that inhibition of Wee1 activity causes gastrulation defects by impairing key morphogenetic movements involved in gastrulation [69].

One of the main roles of the G2/M checkpoint is to assure that no DNA damage exists prior to cell division [17]. Consequently, the cell cycle arrest has been shown to promote cell survival in various cancer types, such as breast cancers, leukemia, melanoma, and adult and pediatric brain tumors [66] and abrogation of this arrest leads to cell death in breast cancer and human cervical cancer [54,55]. In agreement with these evidences, our study demonstrated that inhibition of Wee1 kinase activity resulted in cells progression from G2/M to mitosis, but many of these cells are TUNEL-positive. Thus, the premature termination of the repair process, may have increased the number of TUNEL-positive cells in 12°C-diapaused embryos. Interestingly, it has been previously shown that DNA damage is increased at low temperature in frog tadpole, fish and shrimps [70–73]. DNA damage activates the ATM/ATR DNA damage response mechanisms [74–76], found in tumors and cancerous cells, which in turn induces the expression of *Wee1* [77,78], resulting in cell cycle arrest at the G2/M phase [66]. Moreover, DNA of eukaryotic cells including human cells arrested in the G2/M transition phase has been found to be repaired by homologous recombination [79]. Thus, our result raise the possibility that G2/M arrest of blastodermal cells as a result of induced *Wee1* expression at 12°C could be the response to DNA damage, for better survival of cells in chick embryos diapaused at 12°C [3]. Thus, the evolutionary conserved role of *Wee1*, the similarities between the highly proliferating embryonic cells and related diseases in human patients, may render the chick embryo as a valuable model organism for studying potential therapeutic approaches.

In summary, this study suggests that embryonic survival at 12°C is associated with maintaining blastoderm cytoarchitecture and cell viability as a result of cell cycle regulation by an arrest in G2/M phase mediated by Wee1. In contrast, *Wee1* expression is not up-regulated in embryos diapaused at 18°C, suggesting that the enhanced and abnormal tissue volume at this temperature is due to progression of mitosis. Altogether, our findings provide for the first time a missing link between cytoarchitectural changes, cell proliferation and cell survival mechanisms in diapaused embryos.

## 5. Conclusions

In conclusion, avian embryos must adapt to low temperature exposure at very initial stage of their development, by undergoing diapause. This allows embryos to suspend their development for a long duration and still survive after exiting from it. In this study, we used HREM image analysis to quantify the morphological changes in embryos diapaused at different temperatures, 12°C and 18°C. In particular, we found that the cytoarchitecture of the embryos diapaused at 12°C was better preserved than that of embryos diapaused at 18°C. This characteristic is coupled with an arrest in the cell cycle, at the G2/M stage, by upregulation of *Wee1* at 12°C, but not at 18°C. Inhibition of Wee1 kinase activity in embryos in diapause at 12°C resulted in increased tissue size and higher staining for cell death marker, similar to embryos diapaused at 18°C. By combining HREM imaging technology with RNAseq and molecular manipulation this revealed a new role of Wee1 to safeguard mitotic entry to regulate cytoarchitecture and cell survival in chick embryos during diapause.

## Supplementary Figures

**Figure S1.**
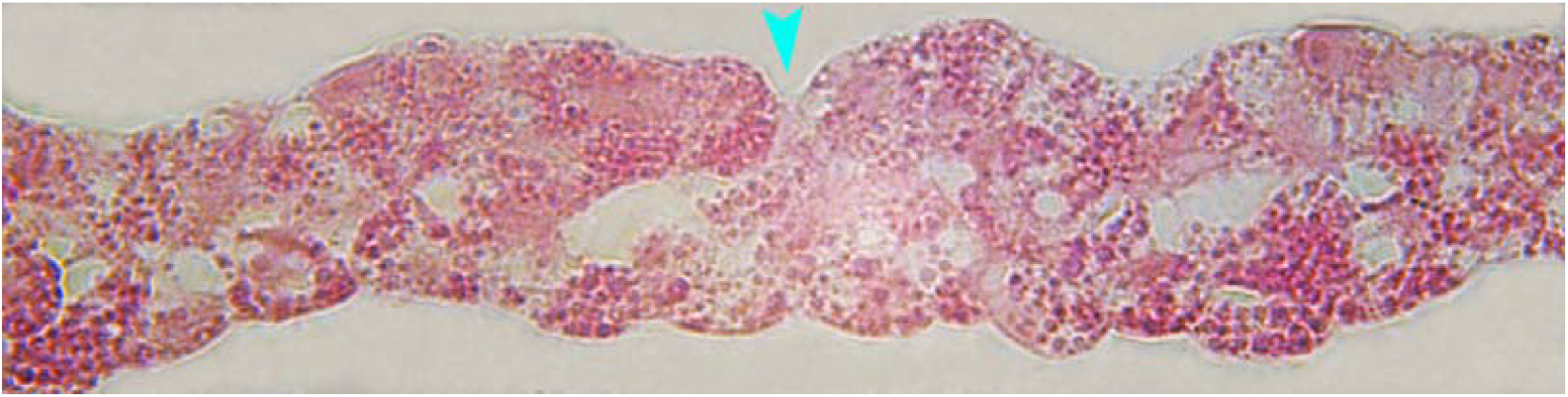
Cellular architecture of blastoderm following prolonged diapause at 18°C. Embryos diapaused at 18°C for 28 d were isolated, fixed in PFA for 24 h, washed with PBS, dehydrated in series of methanol (25%, 50%, 75%, and 100%, each for 20 min) and subjected to HREM analysis as described in the method section. Briefly, embryos soaked with 100% JB4 mix were embedded in plastic molds and accessed for HREM sectioning. The plastic sections were collected and placed on a slide containing a drop of DDW. This allowed the plastic tissue sections to adhere to the slide. The slide was then dried at room temperature and imaged. The captured images were analyzed, and showed thickening of the epiblast regions and formation of recesses in embryos in diapause at 18°C for 28 d (blue arrowhead).

**Figure S2.**
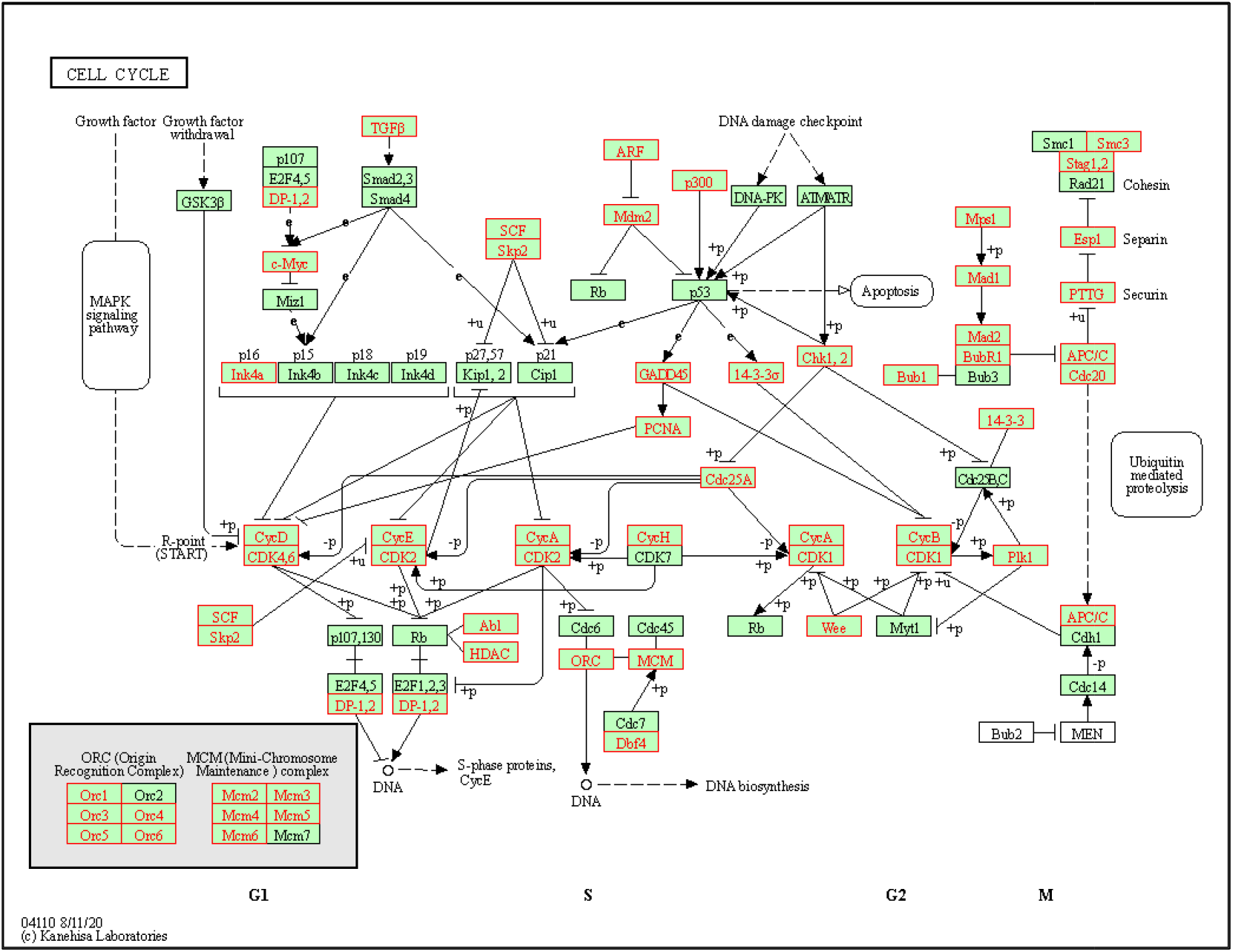
An expanded scheme of enriched cell cycle pathway in 28d/12°C group using KEGG pathway enrichment analysis. To uncover the molecular mechanism responsible for maintaining the cytoarchitecture of the blastoderm following 28 d of diapause at 12°C versus 18°C, RNAseq analysis was performed. Differentially expressed genes between the 28d/12°C and 28d/18°C groups were obtained and genes upregulated in each group were further examined for KEGG pathway enrichment analysis. This showed enrichment of the cell cycle pathway in the 28d/12°C group. The genes are distributed according to the cell cycle phase (G1, S, G2, and M) and enriched gene sets are highlighted in red.

**Figure S3.**
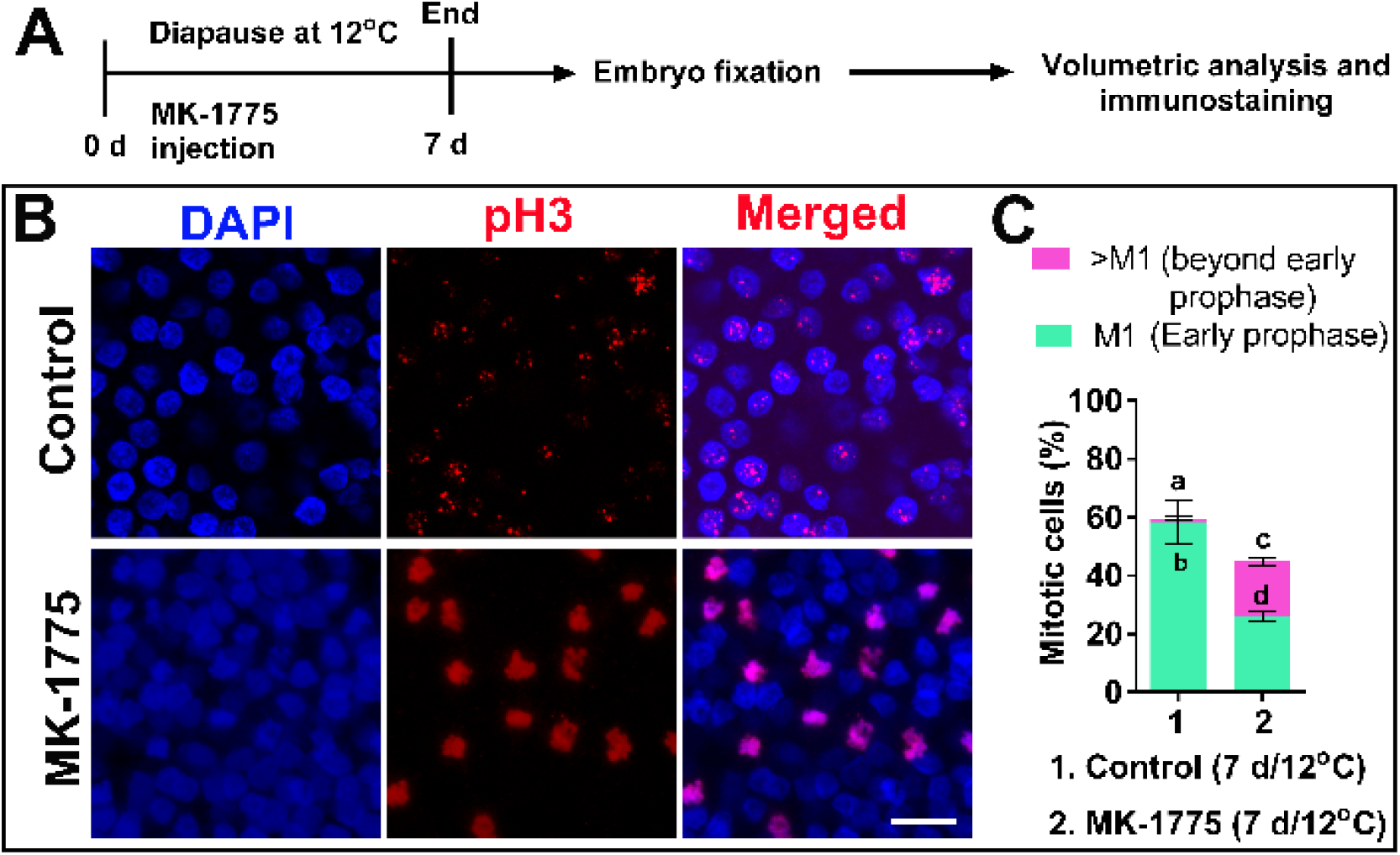
Investigation of the role of Wee1 in regulating the cell cycling of embryos in diapause at 12oC by treating embryos with the Wee1 kinase inhibitor MK-1775. **(A)** Experimental design. Fresh embryos were treated with MK-1775 and subsequently, they underwent diapause for 7 d at 12oC. PBS treated embryos under same condition were used as control. Following treatment, embryos were isolated, fixed, immunostained with anti-pH3 antibody and accessed for confocal microscopy. In a separate experiment, the treated and control embryos underwent volumetric analysis using 3-D image analysis method (Figure 8). **(B)** pH3 immunostaining of embryos shows majority of cells in early prophase in control embryos, whereas, following MK-1775 treatment, cell cycling progresses. Bars 20 *µ*m. **(C)** Quantification of pH3 positive cells following MK-1775 show significantly higher number of cycling beyond early prophase. Different connecting letter refers that the compared groups are significantly different to each other (Two-way ANOVA; a vs. b: p<0.0001; c vs. d: p=0.0207; a vs. c: p<0.0001).

**Supplementary table 1.**
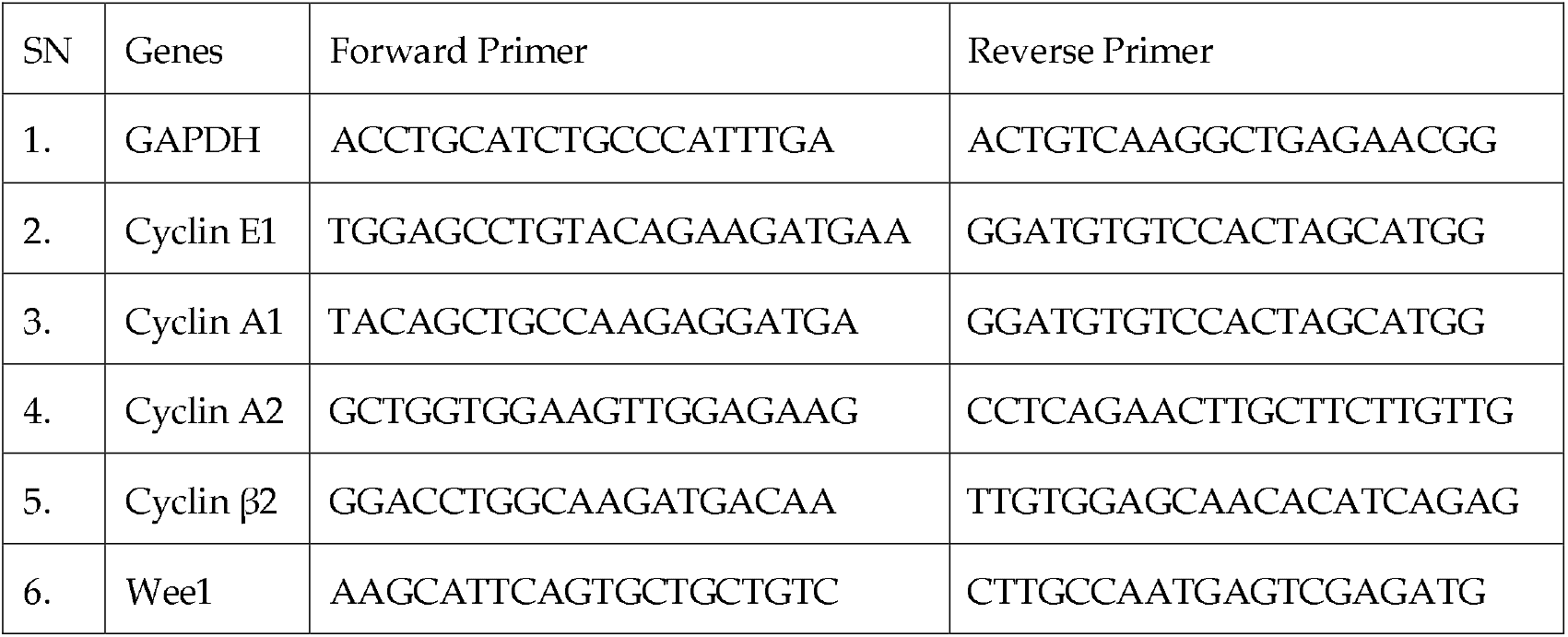
Primer lists

